# System-level analysis of metabolic trade-offs during anaerobic photoheterotrophic growth in *Rhodopseudomonas palustris*

**DOI:** 10.1101/430751

**Authors:** Ali Navid, Yongqin Jiao, Sergio Ernesto Wong, Jennifer Pett-Ridge

## Abstract

**Background:** Living organisms need to allocate their limited resources in a manner that optimizes their overall fitness by simultaneously achieving several different biological objectives. Examination of these biological trade-offs can provide invaluable information regarding the biophysical and biochemical bases behind observed cellular phenotypes. A quantitative knowledge of a cell system’s critical objectives is also needed for engineering of cellular metabolism, where there is interest in mitigating the fitness costs that may result from human manipulation.

**Results:** To study metabolism in photoheterotrophs, we developed and validated a genome-scale model of metabolism in *Rhodopseudomonas palustris*, a metabolically versatile gram-negative purple non-sulfur bacterium capable of growing phototrophically on various carbons sources, including inorganic carbon and aromatic compounds. To quantitatively assess trade-offs among a set of important biological objectives during different metabolic growth modes, we used our new model to conduct an 8-dimensional multi-objective flux analysis of metabolism in *R. palustris*. Our results revealed that phototrophic metabolism in *R. palustris* is a light-limited growth mode under anaerobic conditions, regardless of the available carbon source. Under photoheterotrophic conditions, *R. Palustris* prioritizes the optimization of carbon efficiency, followed by ATP production and biomass production rate, in a Pareto-optimal manner. To achieve maximum carbon fixation, cells appear to divert limited energy resources away from growth and toward CO_2_ fixation, even in presence of excess reduced carbon. We also found that to achieve the theoretical maximum rate of biomass production, anaerobic metabolism requires import of additional compounds (such as protons) to serve as electron acceptors. Finally, we found that production of hydrogen gas, of potential interest as a candidate biofuel, lowers the cellular growth rates under all circumstances.

**Conclusions:** Photoheterotrophic metabolism of *R. palustris* is primarily regulated by the amount of light it can absorb and not the availability of carbon. However, despite carbon’s secondary role as a regulating factor, *R. palustris’* metabolism strives for maximum carbon efficiency, even when this increased efficiency leads to slightly lower growth rates.

## Background

The high-throughput “omics” revolution has resulted in a deluge of system-level information about the components of living organisms. Optimally, integration and interpretation of these data can provide mechanistic insights about cellular behaviors and function. This new information can also be used by synthetic biologists to manipulate the biochemical processes within select (primarily microbial) organisms in order to achieve desired outcomes such as production of valuable compounds like drugs or biofuels. Biofuels generated via microbial metabolism are of significant interest because they could theoretically serve as a primary source of energy for industry and transportation, thus supplanting fossil fuels and mitigating the harmful effects of global climate change.

These positive attributes have resulted in a significant interest in conducting system-level analyses of modes of metabolism that use renewable resources and release part of the generated energy in useful forms. Metabolism in phototrophic organisms, as well as those that can catabolize aromatic compounds (a major component of plant biomass), are such modes of metabolism.

To manipulate cellular metabolism to achieve a desired biological task (or objective), while simultaneously mitigating the fitness costs that may result from human tampering, it is necessary to have quantitative knowledge of the cellular system’s critical objectives. This can ensure that engineering goals do not significantly alter the natural balance of system for a given environment. Life is based on a series of trade-offs that represent the fitness price that organisms pay when improvement in one of their traits results in detrimental change for another; a system state referred to as “Pareto efficient”[1]. Evolution ensures that all living organisms are Pareto efficient; otherwise, on an evolutionary timescale the non-efficient would be outcompeted and outlasted by organisms with better performance in all or some biological tasks.

Knowing the nature and magnitude of biological trade-offs can help to establish the biophysical and biochemical underpinnings of observed cellular phenotypes. This type of knowledge can be gained from genome-scale models (GSMs) and system-level multi-objective analyses of cellular processes. Genome-scale mathematical modeling of metabolic networks is a key tool that has been used in systems biology studies of microbes ranging from model organisms such as *E. coli*[2, 3] and baker’s yeast[4], to those of ecological and industrial interest[5–8], as well as pathogens[9, 10]. A number of models have also been developed for photosynthetic organisms, such as the purple non-sulfur bacterium *Rhodobacter sphaeroides*[11] and the cyanobacterium *Synechococcus* sp. PCC 7002[12]. When used with constraint-based methods like flux balance analysis (FBA)[13, 14], GSMs can quantitatively describe a metabolic network’s fluxes under a steady state assumption. This permits analyses of different types of so-called *‘omics* data via *in silico* simulation of all processes of interest following assorted genetic and environmental perturbations [15].

Despite its many uses[15, 16], the standard FBA approach is insufficient for analysis of trade-offs between large numbers of system objectives. FBA examines the feasible flux patterns in a system while optimizing a single biological objective function. To examine trade-offs between different objectives of a system, a ‘multi-objective flux analysis’ (MOFA) approach is needed. MOFA is based on the widely used multi-objective optimization (MO) method; a critical tool in fields where decision makers need to consider trade-offs between various conflicting objectives. The desired outcomes of MO simulations are called Pareto-optimal (PO) solutions. A PO solution of a problem is one for which any improvement in value of one objective will lead to diminishment of another[17, 18].

To date, a number of important MO analyses of biological processes have been developed using constraint-based models [19–24]. For example, phenotype phase plane analysis is one such method that has been used to study the optimal utilization of a system’s metabolic network as a function of variations of two environmental constraints[25, 26]. Thus, this method examines the interactions between three system objectives (growth and the two constraints). Other MO-based studies have provided important insights for systems bioengineering, including the relationships between environments and regulatory mechanisms [27–29], the minimal number and combination of augmentations to a system that would result in greatest amount of strain optimization [30, 31], and guidelines for tuning synthetic biology devices [32].

Recently Shuetz et al.[33] showed that when examining trade-offs between double and triple combinations of different biological objectives in microbes, a PO combination of three tasks – maximum biomass yield, maximum yield of ATP, and optimal allocation of resources – best explains the measured flux distribution for a variety of organisms and conditions. While these are the top three evolutionarily important objectives and their combined optimization best describes observed metabolic fluxes among all examined trios of objectives, the match is not exact. To improve the match between optimization predictions and flux measurements, Pareto optimization of other biological objectives (pertinent to specific organisms and growth conditions) could help. Examination of other objectives also provides us with quantitative insights into how their activities influence cellular workings.

In this study, we examined energy and carbon trade-offs for different types of phototrophic metabolism. So, in order to gain a more complete understanding of the system, besides the three classical objectives noted above, we examined objectives related to environmental nutritional conditions as well as production of compounds of interest (such as H_2_ gas).

The modeled organism, *Rhodopseudomonas palustris* (RP), is a purple non-sulfur (PNS) proteobacterium from the *Rhodospirillaceae* family. RP’s metabolism is extremely versatile and serves as a model for several important biological phenomena, including biodegradation of industrial waste[34, 35], electricity generation[36], and production of hydrogen gas (H_2_)[37–39]. RP has the capacity to switch between four different types of metabolism (photoautotrophy, photoheterotrophy, chemoautotrophy, and chemoheterotrophy). It can grow in both aerobic and anaerobic conditions while using light and organic compounds as energy sources, and organic or inorganic[39–41] compounds as electron sources. RP can also fix both carbon dioxide (CO_2_) and nitrogen gas (N_2_)[42, 43].

Finally, RP can metabolize aromatic compounds; using them as a carbon source in a light-dependent fashion under anaerobic conditions (LN). New insights gained through our MO system-level analyses of this type of RP metabolism are important for industrial and environmental reasons--since microbial production of biofuels as well as bioremediation of aromatic pollutants may sometimes occur in low oxygen environments.

To determine carbon and energy fluxes among RP’s different metabolic modes, we developed a GSM of metabolism in RP. Although a model of RP’s central carbon metabolism had been previously developed[44], that model did not account for a significant fraction of the metabolic reactions in the system and did not use an RP-specific biomass composition. Our model incorporates most of the metabolic reactions that are catalyzed by enzymes encoded in RP’s genome, as well as orphan reactions that are needed for growth in experimentally tested media. The model also uses an RP-specific biomass composition. Our model has been extensively curated to ensure mass balance, particularly with regards to protons, because as we discovered (and others have noted[45, 46]) poor accounting of protons can result in erroneous outcomes. Specific details of our model building process are described in the methods section.

Although system-level MOFA analyses of metabolism have expanded analyses of biosystems beyond FBA’s canonical single objective optimization, to date, the maximum number of objective trade-offs that have been simultaneously examined has not exceeded eight[28]. For example, in one of the more detailed of these analyses, Schuetz et al. studied trade-offs among a suite of objectives in nine bacteria by examining trade-offs among different combinations of 3–5 objectives [33]. While their approach would work for analysis of a small set of objectives; when analyzing larger sets of objectives, in order to avoid incomplete considerations of feasible functional capabilities of the system, one would need to analyze an ever-increasing number of small subsets of objectives.

Given RP’s versatile metabolism, system-level MOFA analysis of its metabolism necessitated examination of trade-offs among greater than five interdependent biological objectives and environmental constraints. To this end, we developed a computational algorithm for MOFA similar to the one used by Nagrath et al.[20], which uses the Normalized Normal Constraint (NNC) method [47] to generate multidimensional Pareto solutions for our analyses. In our approach, the number of objectives that can be analyzed simultaneously is unlimited. However, as the number of examined objectives increases, we caution that the computational resources needed for the calculations will increase nonlinearly.

In addition to our MOFA analyses of metabolic objective trade-offs, we used the RP GSM to: a) investigate RP’s metabolism of different carbon sources, b) examine the role of proton availability in affecting mode of metabolism, and c) study RP’s capacity to produce H_2_.

## Results & Discussion

We used existing RP flux measurements[48] of photoheterotrophic acetate metabolism to validate our model’s predictions, and constrain our model based on an experimentally observed metabolic phenotype. This allowed us to assess the metabolic limitations of the system. Using this new-found insight and MOFA, we then examined the relative importance of different biological objectives during different forms of mixotrophic metabolism. The results of these analyses are detailed below. We also used FBA to examine the robustness of RP’s metabolism to genetic perturbations (see supplementary materials).

### Metabolic trade-offs during mixotrophic growth

#### 1. Light-anaerobic metabolism of acetate

To quantify the extent to which some biological objectives of RP dictate the behavior of the system under anaerobic mixotrophic conditions, we conducted a system-level MOFA analysis of LN metabolism of acetate[48], assuming that the system operated in a Pareto efficient manner (Fig. 1). LN acetate metabolism was chosen as the test case due to the availability of experimental flux measurements for this mode of metabolism[48]. In order to quantitatively examine how RP allocates its limited resources, we mapped the experimental flux measurements within an 8-dimensional MOFA solution space, including: 1) biomass production (growth), 2) CO_2_ export, 3) ATP production, 4) nutrient allocation (minimal metabolite transport), 5) H_2_ export, 6) pyruvate export, 7) succinate export, and 8) α-ketoglutarate export. The latter three objectives were included to examine the role of carbon fixation as a sink for excess electrons.

**Figure 1.**
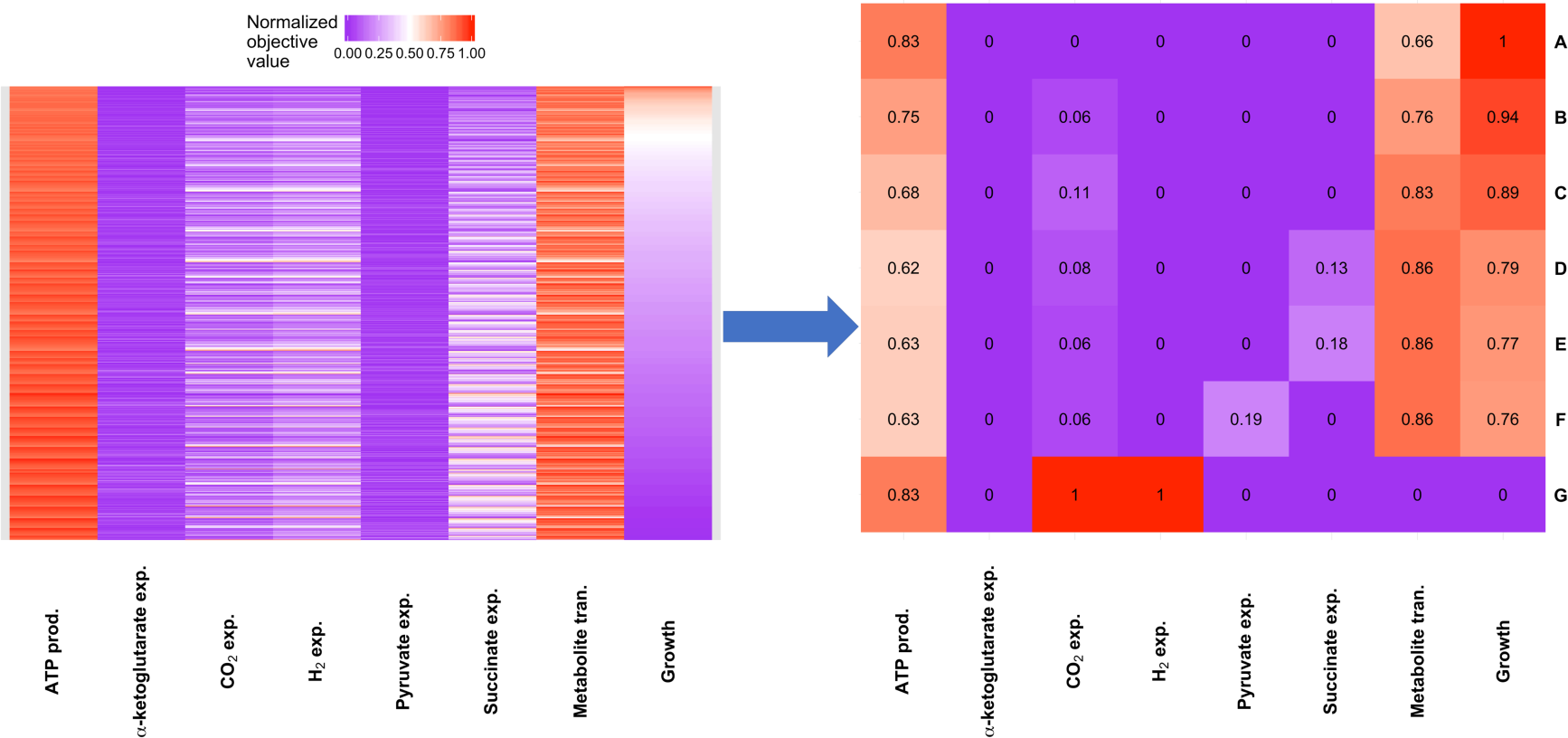
Heat map of the 8-dimensional Pareto front from MOFA analysis (objectives: growth, carbon fixation/carbon efficiency, production of some small organic byproduct (succinate and α-ketoglutarate), and H_2_ production) of anaerobic mixotrophic metabolism of acetate in *R. palustris*. The figure on left displays 1719 unique Pareto-optimal solutions identified during our analysis. In the right panel, select Pareto-optimal solutions from same MOFA study are highlighted for discussion. Each biological objective was examined at intervals equaling 1/5 maximum normalized value. The analyses show that the observed growth rate (E) is smaller than the maximum theoretical growth rate (A) in a carbon-limited system with unlimited light absorbing capability. It also shows that H_2_ production competes with growth pathways for resources and that maximum theoretical H_2_ production (G) would result in cessation of growth. This result is supported by experimental observations[63].

##### Condition 1: Fixed import of acetate

Archetype is a termed coined by Shoval et al. [49], and used by others (e.g. [50–52]), to describe a phenotype that optimizes a single task. When FBA optimizes growth as its sole objective function, it solves for a flux pattern that results in the theoretical growth archetype. For our analyses, we first used FBA to identify the theoretical growth archetype under carbon-limited conditions. At a fixed acetate uptake rate that matched experimental flux measurements (1.96 mmol.gDW^−1^.hr^−1^), the modeled cell did not produce any small carbon byproducts (Row A, Fig. 1). This means that all the CO_2_ generated from acetate consumption was fixed and used for production of biomass. For the theoretical growth archetype, FBA flux predictions indicate that the Calvin-Benson-Bassham (CBB) process fixed the majority of the produced CO_2_ (>70%). Conversion to bicarbonate by carbonate anhydrase (E.C. 4.2.1.1), and pyruvate via pyruvate synthase (E.C. 1.2.7.1), fixed the remaining CO_2_. However, the predicted growth rate for the carbon-limited growth archetype was higher than experimentally measured values (the model predicted a doubling time of 6.4 h vs. measured 8.4 h[53]). This suggests that the amount of carbon imported exceeded the growth demands of RP. Thus, we conclude that under the experimental conditions outlined by McKinlay and Harwood[48], RP’s metabolism was not carbon limited.

To test the significance of carbon fixation toward maximizing cellular growth, we *in silico* inactivated CBB (by fixing the reaction rate for RuBisCO at zero). Elimination of CBB increased the predicted doubling time to 7.2 h, still less than the experimentally measured value (Row C, Fig. 1). In the absence of CBB, the predicted rate of CO_2_ export was 21% of the rate of acetate uptake. This value matches the measured total amount of CO_2_ produced (22% of acetate flux[48]); suggesting that the FBA correctly predicted the LN carbon metabolism pathways.

Given that experimentally observed phenotype includes export of CO_2_, we set the rate of CO_2_ export equal to the measured value (0.23 mmol.gDW^−1^.h^−1^. With unlimited light, the predicted doubling time (6.8 h) was still smaller than the measured value (Row B, Fig. 1). Thus, we found that if light is available and the upper limit for the enzymatic capacity of system to fix carbon has not been reached, CO_2_ would be fixed for systems where maximization of growth is the sole objective of the system.

##### Condition 2: Fixed import of acetate, varying absorption of light

We next examined the effect of light-absorption on the rate of biomass production. Decreasing the amount of light absorbed by RP increased its doubling time, and improved the agreement between model predictions and experimental observations. However, when the rate of light absorption decreased, the rate of CO_2_ export began to rise above experimentally observed values. To examine what caused these observations, we explored two scenarios:

1. At the measured CO_2_ export rate, we reduced the rate of light absorption until the doubling time matched the experimentally measured value. This was achieved at the photon uptake rate of 36.6 mmol.gDW^−1^.h^−1^, suggesting a light-limited metabolism. This is consistent with studies of another phototrophic organism [54], *Synechocystis* sp PCC6803, and also what is known about RP’s primary ecological niche--light-limited environments where it grows beneath cyanobacteria in microbial mats[55]. As we decreased the level of absorbed light, the model predicted that RP begins to export succinate (Row E, Fig. 1; representing 18% of the carbon imported as acetate.
2. When succinate export is blocked, we found a 1% reduction in maximum growth rate and instead pyruvate is exported (Row F, Fig. 1). The reduced growth rate is likely due to higher energy costs associated with the production of pyruvate from acetate, relative to succinate.

##### Condition 3: Fixed light import, fixed CO_2_ export

At a fixed rate of photon absorption, with no limit to acetate uptake, the model predicted that 17% less acetate consumption results in a slightly (1%) higher growth rate. This result suggests that the measured rate of acetate import is higher than the amount needed to achieve the experimentally measured growth rate.

##### Condition 4: Fixed acetate and light import

Finally, the model showed that at a fixed rate of photon and acetate uptake, if the limit on CO_2_ export is removed, CO_2_ export increases, growth rate increases by ~3%, and the rate of small organic acid export drops to ~13% of imported carbon (Row D, Fig. 1). This outcome implies that the measured CO_2_ export rate is lower than what is needed to achieve the theoretically predicted growth archetype. This prediction also implies that the cell uses processes other than CO_2_ export to expunge excess carbon. Instead, the cells divert a fraction of their absorbed and metabolically generated energy towards production and export of small organic acids, likely driven by redox balance or regulatory constraints.

This result slightly differs from what Hadicke et al. found in their *in silico* analysis of redox balancing and biohydrogen production in purple non-sulfur bacteria (PNS)[44]. In their study, Hadicke et al. were able to grow PNS photoheterotrophically by only exporting CO_2_ and biomass[44]. The differences we observe in our analyses are likely because we constrained the model with three measured values (growth, CO_2_ export rate, and acetate import rate) simultaneously and allowed for export of small organic compounds. In contrast, Hadicke et al.[44] blocked the export of all byproducts other than CO_2_ and did not fix the rates of light absorption or CO_2_ export. Thus, they were able to predict photoheterotrophic growth with only CO_2_ and biomass as byproducts (similar to our outcome for condition 1), whereas our model with fixed experimental measurements and light absorption, required export of excess imported carbon via means other than CO_2_.

Mapping the flux measurements[48] for LN growth of RP on acetate within our 8dimensional MOFA-derived phenotype solution space revealed a pattern consistent with Schuetz et al.[33], *i.e*., maximizing efficient resource allocation (97% of optimal value), ATP production (84% of maximum value), and growth (the main FBA assumption, 79% of its theoretical maximum value) are the top 3 biological objectives that are optimized. However, our results also suggest that a fourth objective — production of small organic acids while light energy is available — is also optimized.

Optimization of this fourth objective diminishes the ability of the cell to achieve the growth archetype for the amount of resources that are imported. Model predictions suggest that RP diverts some of its absorbed light energy toward the production of reduced organic compounds. We can gain a better understanding of this process by examining the degree of reduction of various carbon-based compounds imported and exported by the system. The degree of reduction for a compound per carbon atom (κ) can be quantified using the concept developed by Roels[56].

As noted above, our predictions indicate that the cell imports extra carbon and electrons in the form of acetate (κ=4), uses some of this resource for biomass production (κ=4.19) and then uses light energy to export the remaining electrons as organic compounds like succinate (κ=3.5), that although more oxidized than acetate are significantly more reduced than CO_2_ (κ=0). One possible explanation for this behavior could be that the cell stores some of the available energy as easily metabolized compounds for consumption during dark periods, a behavior that has been observed in cyanobacteria[57]. However, the amount of light and carbon the model predicts to be used for the production of reduced byproducts is small and well within the range of uncertainties of experimental measurements. Sensitive experimental analyses to identify and quantify the metabolic byproducts of LN metabolism could either validate our MOFA prediction of organic carbon export or support the previous assumption that CO_2_ is the sole byproduct[44].

### 2. Differences between RP’s metabolism of aliphatic and aromatic carbon sources

Our result for aliphatic metabolism of RP, and previous analyses[44], highlights the importance of carbon fixation for achieving the growth archetype in phototrophs like RP. Interestingly, transcriptomic measurements suggest that RP upregulates genes associated with CBB after switching from growth on an aliphatic carbon source (succinate) to aromatic compounds like 4-coumarate or benzoate[58]. Given our prediction that CBB is the primary pathway of carbon fixation during aliphatic metabolism in RP, the increased use of CBB for aromatic metabolism warranted further investigation. To this end, we used GX-FBA[59], an *in silico* method that uses transcriptomic data to constrain GSMs (see methods), and makes it possible to examine metabolic changes as an organism transitions from one environment to another.

We used previously-collected transcriptomic data[58] to explore RP’s metabolism as it switches from aromatic to aliphatic carbon sources. Figure 2 illustrates changes in metabolic pathway fluxes as RP’s carbon source shifted from 4-coumarate to succinate. GX-FBA predicts that this transition results in a reduction of CBB activity, consistent with measurements that suggest downregulation of genes associated with this pathway[58]. We (and others[58]) attribute this to the fact that the examined aromatic compounds are more reduced (κ>4. 1) than succinate (κ=3.5) and hence need greater use of CBB as an electron sink. In addition, GX-FBA predicted that switching to an aliphatic carbon source results in an increase in fluxes through cysteine (κ=5.67), methionine (κ=6), and pyruvate metabolic pathways. Given the highly reduced nature of these sulfur-based amino acids, it is curious that a switch from the more-reduced aromatic to less-reduced aliphatic carbon source results in increased production of these compounds, and indicates that there is a reason beyond redox balance associated with these metabolic changes.

**Figure 2.**
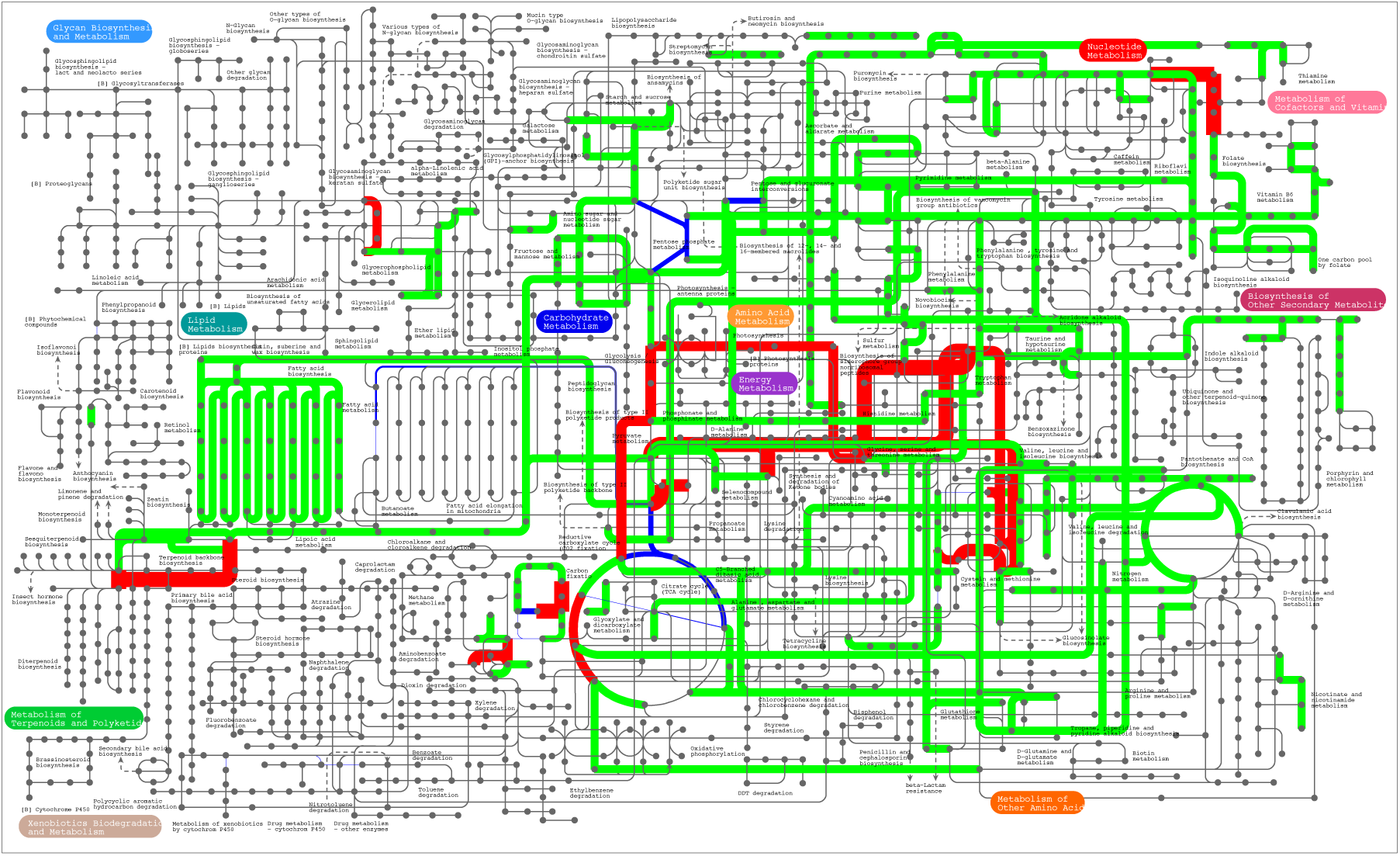
GX-FBA predicted change in metabolic pathway activity in *Rhodopseudomonas palustris* after changing the carbon source from 4-coumarate to succinate. The transition leads to reduced carbon fixation via CBB. Blue=flux decrease, red=flux increase, green=flux did not increase or decrease by at least a factor of 2. The graph is made using the iPath2 program[115] and the width of the lines (w) is set to: 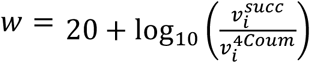. If the calculated w≤0 for sake of being able to notice the change w=1.

The aromatic to aliphatic change in carbon source also results in a decrease in fatty acid (κ>4.67) metabolism, metabolism of lysine (κ=4.67) and tryptophan (κ=4.18), and activity of the benzoate degradation pathway. The reduced activity of the benzoate pathway would be expected and could be linked to reduced production of the aromatic amino acids. Reduced availability of electrons following the switch could explain why production of highly reduced fatty acid compounds is predicted.

### 3. Light-anaerobic metabolism of aromatic compounds

We conducted MOFA analyses to examine the metabolic trade-offs of different biological objectives for RP consuming various aromatic compounds (Table 1). Under light-limited anaerobic conditions with 4-coumarate as the sole carbon source, a small fraction (4.6%) of the imported carbon was exported as CO_2_, while the majority of the produced CO_2_ (83%) was incorporated into biomass. Unexpectedly, at the growth archetype for this environment, our model predicted that the CBB cycle was not the primary route of carbon fixation. Instead, a large fraction (~61%) of CO_2_ was fixed by the enzymes pyruvate:ferredoxin oxidoreductase (E.C. 1.2.7.1) and 2-oxoglutarate synthase (E.C. 1.2.7.3). Both of these enzymes are associated with the reductive citric acid cycle (rTCA) which is a pathway for carbon fixation in some photoautotrophic organisms such as green-sulfur bacterium *Chlorobium limicola*[60] and chemolithoautotrophic archaea[61].

**Table 1.** Summary of model predicted characteristics of light anaerobic mixotrophic metabolism of three aromatic compounds. In each case, the model predicted doubling time is smaller than the measured value. To achieve the theoretical maximum growth rates, the cell must extensively use rTCA (a carbon inefficient pathway) to fix CO_2_.

| Aromatic substrate | Predicted minimum doubling time (hours) | Measured doubling time (hours)[64] | Rate of substrate import (mmol/gDW.hr) | Percent of imported carbon as CO<sub>2</sub> |
| --- | --- | --- | --- | --- |
| 4-Coumarate | 9 | 9.4 | 0.33 | 4.6 |
| 4HBZ | 8.8 | 12 | 0.44 | 8 |
| Benzoate | 8.7 | 9.3 | 0.43 | 3 |

The presence of rTCA in RP is intriguing since it is not present in another well-studied, closely related species, *Rhodobacter sphaeroides*. One advantage of using rTCA for carbon fixation is that it requires less energy than CBB[61]. Thus, given the energy-limited state of RP cells when light absorption is restricted, using this mode of carbon fixation may be essential for achieving the growth archetype for this condition.

However, the rTCA-based optimum theoretical growth rate for RP was 4% higher than the measured value. While this difference is well within the range of experimental variation for measuring doubling times, we still examined MOFA results at the growth rate that matches experimental measurements (Fig. 3, F). At this lower growth rate, the system can use CBB for carbon fixation (accounting for ~56% of the produced CO2). The switch to CBB also resulted in a 5% improvement in carbon utilization efficiency. Gene expression analyses have verified extensive use of CBB during LN metabolism of 4-coumarate[58]. Thus, our results indicate that under light-limited conditions, RP does not achieve the theoretical growth archetype and instead the primary objective during LN metabolism of 4-coumarate is maximum growth while striving to achieve maximum metabolic efficiency.

**Figure 3.**
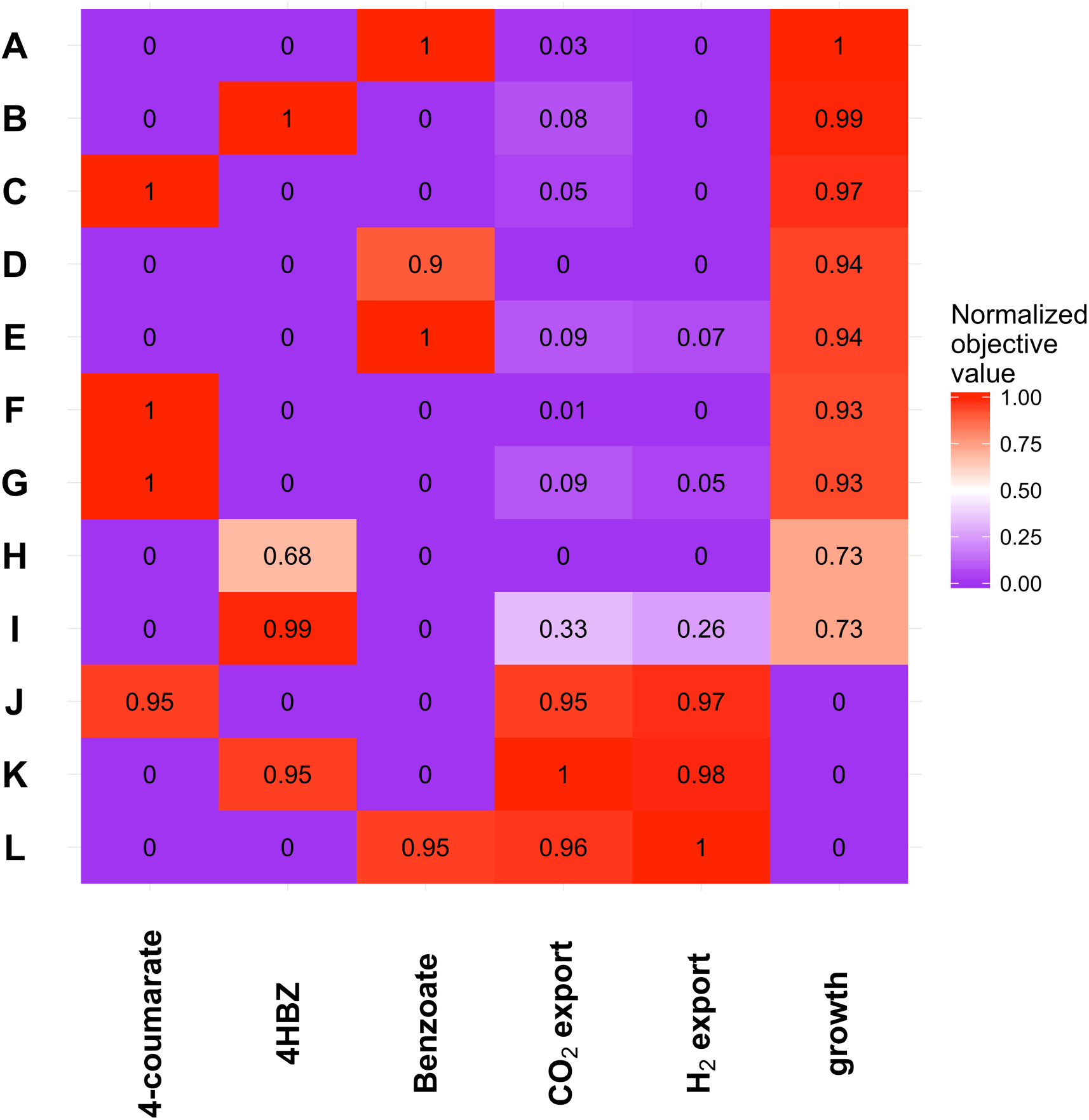
Select Pareto-optimal solutions from the MOFA analysis of growth, carbon fixation/carbon efficiency and H_2_ production in RP when growing on a variety of different aromatic compounds. The analyses show that although the observed metabolisms of different compounds (D, E & F) are lower than the predicted maximum growth rates, they use lower amounts of carbon and fix greater amounts of CO_2_. As with aliphatic metabolism, maximum production of H_2_ results in cessation of growth and full oxidation of the carbon source (G, H & I). The complete set of Pareto solutions are available in the supplementary materials (Figure S1).

Similar analyses of growth archetype with benzoate as the sole carbon source, under light-limited anaerobic conditions (Fig. 3, A), predicted that only 3% of the imported carbon was exported as CO_2_ while approximately 90% of the produced CO_2_ was fixed into biomass. However, as we found for 4-coumarate, due to energy considerations, 67% of CO_2_ was fixed by rTCA. MOFA results at the experimental measured growth rate (Fig. 3, D) showed that the system can switch to CBB for CO_2_ fixation (~50% of generated CO_2_). This mode of metabolism is about 2% more carbon efficient than the metabolism at the optimum theoretical growth rate.

We used the model to examine flux patterns in the growth archetype when RP consumes 4-hydroxybenzoate (4HBZ) under light-limited anaerobic conditions (Fig. 3, B), and found that 8% of the imported carbon was exported as CO_2_. Only about half of the CO_2_ that was produced was fixed via activity of rTCA, while around 20% was fixed through formation of carbonic acid. The results at the measured growth rate (which was significantly smaller than the predicted growth archetype value) identify a likely change in the mode of carbon fixation. At the measured growth rate (Fig. 3, H), the carbon efficient CBB pathway can become the primary route of CO_2_ fixation (83% of produced CO_2_). This mode of metabolism is 8% more carbon efficient than the one used to achieve the theoretical optimum growth rate.

Overall, when simulating light-limited (absorption values similar to those calculated for acetate metabolism, 36.6 mmol.gDW^−1^.h^−1^) photoheterotrophic growth of RP on aromatic compounds, with exception of metabolism on 4HBZ, the model predicts growth rates that are reasonably close to measured values. This can be viewed as further proof of light-limited nature of RP’s phototrophic metabolism.

Also, if one assumes that RP’s enzymatic capacity to fix CO_2_ by rTCA is comparable to that of CBB, then it appears that optimum growth is not the sole objective that controls LN metabolism of aromatics in RP. Our results indicate that the cell grows at the maximum growth rate that also optimizes carbon efficiency. Hence, the cell uses the more energy-expensive CBB carbon fixation pathway, which results in a lower growth rate relative to the theoretical growth archetype, but minimizes carbon waste.

### 4. Proton economy of light-anaerobic metabolism

Our simulations of RP acetate metabolism under LN conditions indicate that RP must import protons from the surrounding medium in order to achieve the measured growth rate. This is congruent with previous studies that have showed exchange of protons with the growth medium is important for maximizing cellular growth[2]. It also has been shown that in other species of *Rhodopseudomonas*, lower pH values in the surrounding medium result in increased rate of biomass production[62].

Compared to biomass, acetate has a higher oxygen to carbon atoms ratio (acetate=1, biomass=0.54). The model predicts that during RP’s LN acetate metabolism, the imported protons bind to excess oxygen atoms of acetate and are exported as water; and without proton uptake, the growth rate is 13% lower (9.7 h doubling time) due to excess oxygen being exported as α-ketoglutarate, wasting carbon and electrons that could otherwise be used for growth. α-ketoglutarate is exported because it has a low degree of reduction (κ=3.2). If we eliminate export of α-ketoglutarate, this further reduces the growth rate (10.5 h doubling time), presumably because alternative oxygen carriers (*e.g*., pyruvate and succinate (κ=3.5)) contain more reduced carbon than α-ketoglutarate. Our model’s prediction that RP imports protons during LN acetate metabolism suggests that an increase in pH of should occur. This consistent with our experimental observations; phototrophic growth of RP on acetate in poorly buffered minimal media leads to a medium pH increase (from 6.7 to 7.2) (Fig. 4).

**Figure 4.**
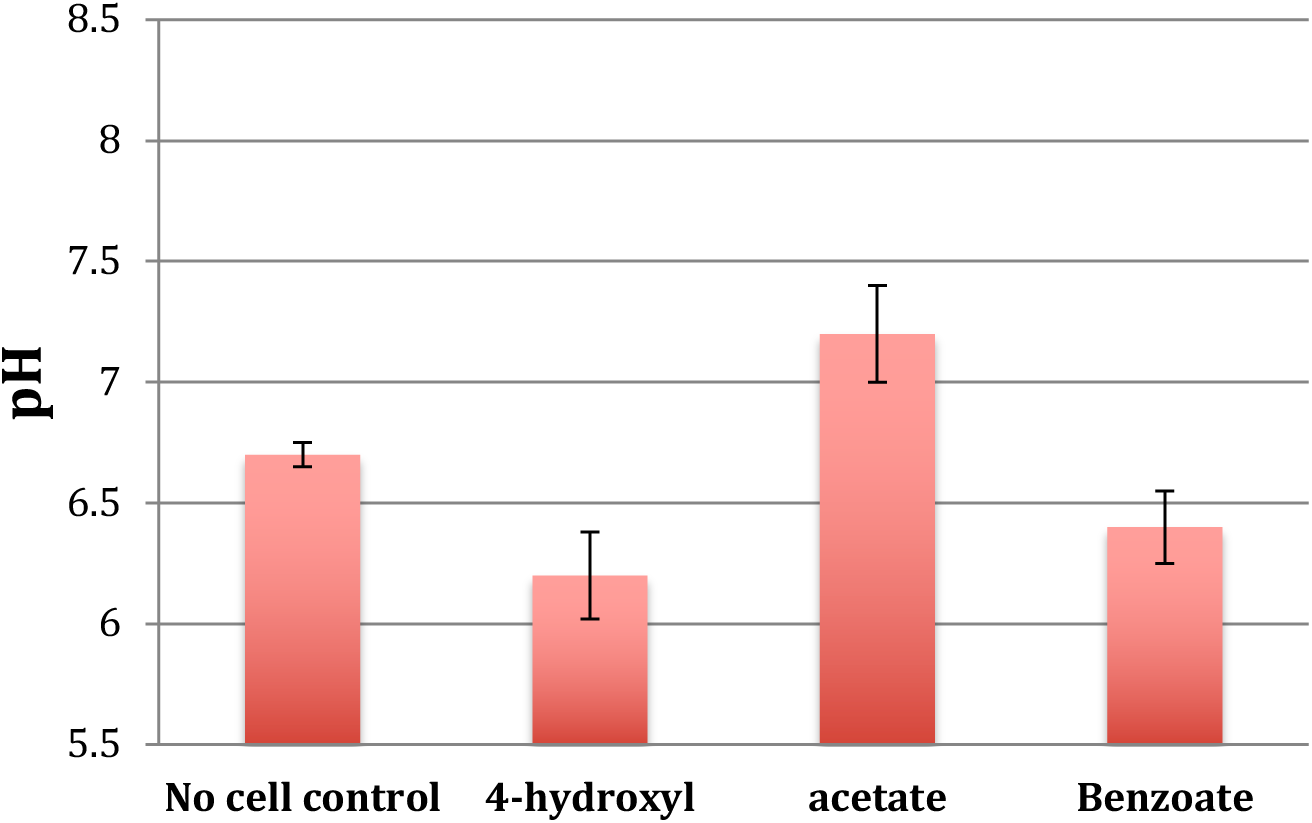
Experimentally measured changes in pH of the growth medium following anaerobic metabolism of 4-hydroxybenzoate, acetate, and benzoate.

Examining the proton economy of LN metabolism of aromatic compounds, the model predicts that unlike metabolism of acetate, breakdown of some compounds *(e.g*., benzoate and 4-coumarate) result in production of protons that are exported. This is due to the low (compared to biomass) hydrogen and oxygen content of these aromatic compounds. The reactions for metabolism of 4-coumarate and benzoate are:

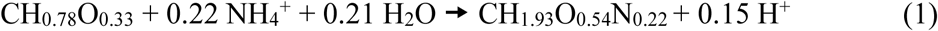

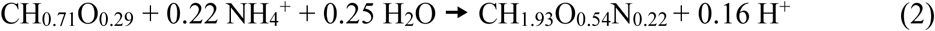

Each carbon that is imported into the cell carries fewer protons and oxygen atoms than acetate (CH_1.5_O) and thus water needs to be used to make up for this shortcoming. Our experiments verified that LN metabolism of benzoate indeed reduces the pH of the growth medium (Fig. 4).

Under light limited conditions, at the measured growth rate, the model predicts that metabolism of 4HBZ should result in import of protons from the medium. This is because the amount of water imported satisfies the oxygen difference between 4HBZ and biomass but not the hydrogen difference. The 4HBZ metabolism equation is:

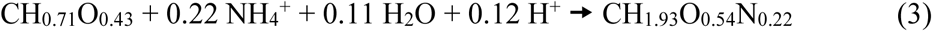

However, our laboratory experiments indicate that LN metabolism of 4HBZ reduces the pH of the medium (Fig. 4). We attribute this discrepancy to the fact that the model-predicted metabolism utilizes the theoretical minimum amount of carbon necessary to achieve the growth archetype. If the 4HBZ metabolism of RP is any less carbon efficient than the model prediction, then the proton metabolism of the system would change. This is because production and export of CO_2_ would increase the amount of H_2_O that needs to be imported to maintain the elemental balance of oxygen. Breakdown of water would result in greater production of protons. For example, if like acetate[48] 10% of the imported carbon is exported as CO_2_, the metabolic process becomes proton producing and the metabolic equation becomes:

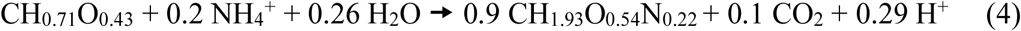

Yet as with acetate, when we compared our experimental observations with the model, it appeared that the behavior of the cell did not solely optimize a single biological objective--such as growth or maximum nutrient use efficiency--but rather a combination of multiple objectives.

### 5. Hydrogen gas production by R. palustris

Converting solar energy into H_2_ gas, a clean-burning alternative fuel, is an environmentally sound means of replacing the use of polluting and finite fossil fuels. RP is a model organism capable of phototrophic production of H_2_. As part of our analysis of RP’s metabolism, we were interested to assess if it is possible to improve its H_2_ production with minimal perturbations of the normal workings of the cell. Examining hydrogen gas production that resulted from *R. palustris’* LN metabolism of acetate, our model predicted a maximum H_2_ production yield of 4 moles H_2_/mole acetate, matching the previously published value[44]. The model also suggests that at both theoretical maximum and observed growth rates, RP should not produce H_2_ gas (Rows A and E, Fig. 1), in agreement with previous experimental observations[48]. MOFA results showed that H_2_ production negatively effects RP’s growth rate and carbon efficiency (*i.e*., carbon fixation via CBB pathway) (Fig. 1, G). While production of small amounts of H2 was necessary for the production of small organic acids like pyruvate and α-ketoglutarate, only CO_2_ production was positively affected by H_2_ production. In solely carbon limited conditions, achieving the maximum theoretical H_2_ production required that RP fully oxidize acetate to CO_2_ and use all the energy generated from this process, as well as photon absorption, to produce H_2_ gas (Row G, Fig. 1).

Simulating nitrogen starvation while growing on acetate, our model predicts that RP will produce the subunits (C_4_H_6_O_2_) of polyhydroxybutyrate (PHB), if this compound is allowed to be exported as a metabolic byproduct. When we optimized PHB production while minimizing the total exchange of nutrients and byproducts (at fixed rates of carbon and photon import), RP produced water, PHB (1 PHB/6 acetate), and succinate. Previous studies have shown that nitrogen starved, non-growing RP cells produce H_2_ gas along with PHB and α-ketoglutarate and CO_2_ as metabolic byproducts[63]. To test whether we could predict the same metabolic phenotype for the nitrogen-starved and non-growing condition, we blocked export of succinate. This constraint resulted in export of α-ketoglutarate as a byproduct. We also observed that while the total amount of PHB produced was lower than when succinate was exuded as a byproduct, the efficiency of PHB production increased (1 PHB/5 acetate). Thus, it appears that nitrogen-starved RP cells simultaneously maximize PHB production and carbon efficiency while minimizing transport fluxes.

MOFA analyses of H_2_ production during LN metabolism of three aromatic compounds indicate that regardless of carbon source, at the theoretical growth archetypes, RP would not produce H_2_ gas (Fig.3, A, B, C). Although the examined aromatic compounds (average κ=4.17) are more reduced than acetate (κ=4) and usually carry extra protons (Equations 1 & 2); at the growth archetypes, the available energy and reducing equivalents are used to fix CO_2_. When RP grows at the experimentally measured rates (which are lower than the predicted growth archetype rates), cells can use excess energy that is not used for production of biomass to produce H_2_.

Production of H_2_ can be induced if the objective of RP’s metabolism is changed from maximized carbon efficiency to one where extra carbon is imported and wasted as CO_2_. For example, when growing on 4-coumarate at the measured growth rate, the model predicts that RP can produce 1.1 mole of H_2_ for every mole of 4-coumarate metabolized (Row G, Fig. 3). This number is similar to that for benzoate (1.14 H_2_/benzoate) (Row E, Fig. 3), both of which are significantly smaller than the ratio for 4HBZ (4.6 H_2_/4HBZ, Row I, Fig. 3). This significantly higher ability to produce H_2_ while metabolizing 4HBZ is due to the fact that the measured growth rate for 4HBZ is ~25% slower than for benzoate and 4-coumarate[64]. Thus, in theory, to produce 5 molecules of H_2_ per 4HBZ, the cell imports 45% more carbon and does not use the extra energy available from absorbing light to fix CO_2_ via CBB. The available excess energy is instead used to produce H_2_ gas. However, as with acetate metabolism, for maximum H_2_ production, MOFA analyses predicted absolute cessation of growth (Rows J, K, L, Fig. 3).

Overall, when we examined H_2_ production via phototrophic metabolism of RP, our results consistently showed that production of H_2_ diminishes the activity of other important biological objectives such as optimum growth or metabolic efficiency (Figs. 1 & 3). This is consistent with previous proposals that H_2_ production competes with biomass generation for resources such as energy, protons, and electrons[63, 65]. Thus, while H_2_ production can serve as an electron sink similar to CBB; the important difference between them is that the latter conserves cellular resources while the former (due to absence of uptake hydrogenase activity[66] in RP) wastes it.

It appears that expression of nitrogenase automatically results in H_2_ production as long as the redox state of the system provides the needed reducing agents[63, 67]. To lower the cost of nitrogenase activity, RP regulates nitrogenase expression through nitrogen sensing and post-translational modification[68]. We expect that under normal (i.e., not nitrogen-limited) conditions, nitrogenase activity is extremely deleterious to cellular growth as well as a number of other cellular objectives. The enzyme’s main function is to fix nitrogen under limiting conditions, and given the importance of this task, we hypothesize that it has a very high affinity for its essential substrates, namely reduced ferredoxin and protons. In support of this, computational analyses have shown that the active site of the nitrogenase enzyme has more affinity for protons and electrons than the platinum-based catalysts that are used for abiotic production of H_2_[69]. Hence, if nitrogenase is expressed, irrespective of the primary objective of the cellular metabolism, it will siphon reduced ferredoxin and protons from other important metabolic processes and (depending on the redox state of the system) produce H_2_.

However, these resources are essential for a variety of other important functions. This could explain why layers of transcriptional control (e.g., nifA[70]) and post-translational regulation exist to tightly control nitrogenase activity.

## Conclusions

We used a multi-objective metabolic flux analysis approach, MOFA, to describe phototrophic metabolism of *Rhodopseudomonas palustris*; simultaneously examining more cellular objectives than any previous metabolic flux analysis study. Our results indicate that the rate of light absorption limits cellular growth. While our analyses indicate that RP primarily optimizes growth, ATP production, and metabolic efficiency, they also suggest that RP’s phototrophic metabolism is energy limited and this has defined the order of importance of these objectives. Our results show that during LN phototrophic metabolism in RP, optimum allocation of resources and ATP production are more important than growth. Our results also hint at a preference for a fourth cellular objective during phototrophic growth, *i.e*., production and excretion of reduced carbon compounds that may be used as an energy source during dark periods. We also found that proton metabolism plays a key role in shaping the observed growth phenotypes. Under anaerobic conditions, the ratio of carbon, oxygen and hydrogen of RP’s carbon source in comparison to its biomass, and the overall carbon efficiency of phototrophic metabolism, determines whether the system prefers a more basic or acidic medium for growth.

## Methods

### Metabolic network reconstruction

Our metabolic network reconstruction for *Rhodopseudomonas palustris* (model iAN1128) is based on the annotated genome of *Rhodopseudomas palustris* CGA009[42]. Of the 4836 predicted genes present in the genome, 1514 are related to cellular metabolism and biosynthesis. Our model accounts for the activity of 1128 of these genes (75%), resulting in 1000 enzymatic reactions. Additional literature surveys identified the activity of 37 local orphan enzymes (13-critical for biomass production, 20-based on literature, 4-pathway hole-filling) and 14 non-enzymatic reactions, resulting in a final model of 1037 reactions and 949 metabolites.

In cases where the roles of essential regulatory genes were known, such as the need for hbaR (RPA0673) and aadR (RPA4234) for growth with 4-hydroxybenzoate as the carbon source[71, 72], these associations were incorporated in the model’s gene-protein-reaction (GPR) basis. But for situations where deletion of a gene reduces RP’s growth rate only under specific media conditions (*e.g*., badR (RPA0655) mutants grow slowly on benzoate[73]), then the gene was not included in the GPR.

The biomass equation for RP was developed using a variety of data sources. The overall breakdown of cellular components is drawn from McKinlay and Harwood[48]. The amino acid, nucleotide, cofactors, carotenoids and phospholipid composition of the biomass are unique to RP. It has been shown that the composition of RP’s cellular membrane changes when the cell transitions between dark-aerobic environments to LN environments[74, 75]. We implemented this change in our model by developing two separate biomass compositions, with bacteriochlorophyll composition of LN biomass drawn from Firsow et al[76] and composition of lipids and fatty acids (both dark and light conditions) drawn from Wood et al[74]. The composition of the polysaccharide moiety of lipopolysaccharides is from Weckesser and Drews[77]. Although in most photosynthetic organisms, genes for carotenoid biosynthesis are simultaneously expressed with other genes involved in chlorophyll biosynthesis and light harvesting process[78], and overall carotenoid concentrations greatly increase between dark and light conditions[79], we did not remove carotenoids from the biomass composition in dark aerobic conditions. This is because carotenoids have other functions in dark conditions, such as quenching free radicals and roles in overall cellular response to environmental stress [80].

It is known that oxygen is not required for the oxidative reactions that are involved in biosynthesis of carotenoids, different forms of quinones, nicotinates, and nicotinamides[81–83]. However, the enzymes associated with these anaerobic transformations are not known. In our RP model, anoxygenic reactions for production of these compounds were drawn from the Model SEED database[84], and were used as orphan reactions without any GPR association.

One significant challenge encountered during the course of our RP GSM development was the unique structure of RP’s Lipid A. The lipid A base of lipopolysaccharides (LPS) in RP has been shown to be composed of 2,3-diamino-2,3-dideoxyglucose[85]; however, the metabolic pathway for production of this compound is unknown. We used microarray analyses to measure the expression of genes known to be associated with usual pathways of LPS production to test whether common pathways for glucosamine-based lipid A synthesis were active in RP. Our analyses showed that most of these genes were prominently expressed in RP. Given this information, we used a number of *in silico* methods such as AS2TS[86] protein structure modeling tool and tools for identification of catalytic sites[87] and protein function predictions (CATSID)[88] to assess whether any of these enzymes could catalyze production of diamino-glucose. However, based on our analyses of RP proteins, we could not find any enzyme able to catalyze the required chemical reactions.

Our model’s biomass has an elemental composition of CH_1.93_O_0.54_N_0.22_. This formula is somewhat different from that measured for the elemental composition of RP strain 42OL (CH_1.8_O_0.38_N_0.18_)[89]. However, the model’s biomass is closer in composition and degree of reduction per carbon mole (κ=4.19) to the “standard” biomass formula of CH_1.8_O_0.5_N_0.2_[90] (κ=4.2) than the composition for strain 42OL (κ=4.5). Hence for our simulations the overall formula for conversion of acetate to biomass is:

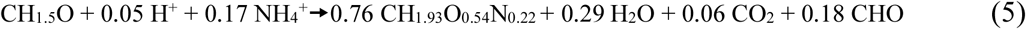

We set the value for non-growth associated maintenance ATP usage to that previously used for *Escherichia coli* (7.6 mmol/gDW h^−1^)[91]. Variation of this value does not change the outcome of metabolic simulations since changing the rate of light absorption will account for any increase or decrease in this value.

We curated the model extensively to ensure absolute mass balance, including proper proton balance under physiological pH values. We also imposed the loop law on the model and eliminated all thermodynamically infeasible type III extreme pathways[92].

For the preprint, the model is included as an excel file with the supplementary materials. Please contact the corresponding author for an SBML version of the model. An SBML file of the model will be included with the Supplementary Material upon publication. At that time, the model can be also downloaded from bbs.llnl.gov/AliNavid.html and EMBL-EBI’s Biomodels Database[93].

Comparing the predicted metabolic phenotypes with experimental observations validated the model. We examined the model’s ability to consume a variety of different carbon sources as reported in the literature[40, 94]. It is known that strains of RP can consume a large array of different aromatic compounds[95–101]. However, while the mechanism for breakdown of the common intermediate in the process of anoxic aromatic catabolism (i.e., benzoyl-coa) [71, 102–107] has been extensively examined, the enzymatic process and associated genes for breakdown of some parent compounds are not known. Furthermore, strain CGA009 cannot consume some of the aromatic compounds that other strains catabolize. For example, while strain CGA009 cannot use 3-chlorobenzoate[108], RP strain RCB100 uses this compound as a carbon source[109]. Thus, our model only accounts for metabolisms of aromatic compounds whose degradation pathways have been identified (benzoate, 4-hydroxybenzoate, phenol, cresol, coumarate, protocatechoate, vanillate, phenol, and cinnamate).

### Flux Balance Analysis

The FBA modeling approach uses a genome-scale metabolic reconstruction as its basis. The reconstruction is developed using elementary functional information derived from annotated genomes and available knowledge of enzymology. The reconstruction of an organism’s metabolic reactions is represented as a stoichiometric matrix, *S(m×n)*, where *m* is the number of metabolites and *n* the number of different reactions. Applying the assumptions of mass balance and metabolic steady-state, the following set of linear equations govern the system’s behavior:

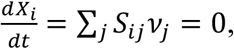

where *X_i_* is the concentration of metabolite *i*. For FBA, other limitations are imposed on the system based on experimental studies, including a limit on the amount of flux that courses through a reaction, as well as constraints on the amount of nutrients imported, and the waste products secreted from the system. These constraints are formulated as:

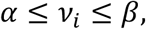

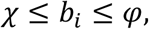

where *b_i_* and *ν_i_* are the export/import flux of metabolite species *i*, and the flux through internal reaction *i* respectively, and *α*, *β, χ*, and *φ* are the lower and upper limits for these fluxes. Finally, FBA utilizes linear programming to determine a feasible steady-state flux vector that optimizes an objective function, most commonly chosen to be the production of biomass, *i.e*. cellular growth. FBA was used with our RP GSM to analyze single gene knockout phenotypes for all the genes in the model. Several reviews[110, 111] provide detailed description of this process.

### GX-FBA

In order to assess differences in RP metabolism when growing on aliphatic and aromatic carbon sources, we used the GX-FBA modeling methodology[59] with available gene-expression measurements[58] for RP growing on different carbon sources. We combined mRNA expression data with a constraint-based framework using the multi-step approach previously detailed for GX-FBA[59]. Note that, for our analyses we chose to only take into account gene-expression changes of at least 50% (±0.5 fold change).

A brief description of GX-FBA steps is:
1. Generate the flux distribution 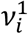 for the starting condition (1) using an Interior Point optimization algorithm with biomass growth or any other appropriate goal as the objective function.
2. For nutritional constraints associated with the post-transition environment (condition (2)), flux variability analysis (FVA)[112] with minimal flux for biomass production set to zero is utilized to calculate the lower and upper fluxes that each model reaction *i* (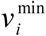 and 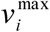 respectively) can carry solely based on environmental limitations and network connectivity. From these results, the mean possible flux value for each reaction 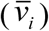 and average flux carried by all active reactions 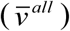 is determined.
3. Identify the set of reactions *T* for which an mRNA expression value can be associated. For protein complexes and reactions catalyzed by isozymes, the maximal up- or down-regulation value is used unless the mRNA expression values are inconsistent (mixture of up- and down-regulation). In the latter case, the reaction is excluded from *T*.
4. Construct a new objective function:

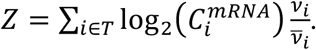 If the flux value for condition 1 of a reaction *i* is zero, 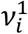 and 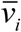 are set equal to the average value for all active reactions 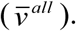. For a more detailed description of this method see Navid and Almaas (2012)[59].

### Catalytic site identification server

The catalytic site identification (CATSID) web server[87, 88] rapidly identifies structural matches to a user-specified catalytic site among all Protein Data Bank proteins. It also examines a user-specified protein structure or model to identify structural matches to a library of catalytic sites. CATSID includes a database of pre-calculated matches between all Protein Data Bank proteins and the library of catalytic sites. The databank has been used to derive a set of theorized new enzymatic function annotations. Matches and predicted binding sites can be visualized interactively online. We used CATSID along with a number of other *in silico* methods for examining protein structure such as AS2TS[86] to determine if whether any of the enzymes encoded by RP genome could catalyze production of diamino-glucose, a key subunit of RP Lipid A.

### Multi-objective Flux analysis

As with the effort by Nagrath et al.[20], our MOFA program uses the Normalized Normal Constraint (NNC) method[47] to map the *n*-dimensional Pareto front of the competing metabolic objectives. NNC generates an even distribution of Pareto solutions on convex or non-convex Pareto frontiers for problems of *n*-objectives. Additionally, NNC is usable for an arbitrary number of objectives and its results are entirely independent of the scales of the examined objectives scales.

The mathematical representation of a generic multi-objective optimization problem is:

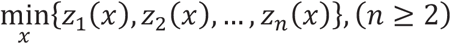

subject to:

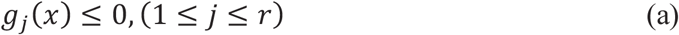

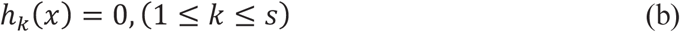

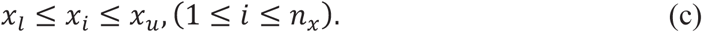

Vector *x* denotes the design variables (fluxes) and *z_n_* denotes the *n*th objective function. Equations a-c denote the inequality, equality and side constraints.

NNC can be briefly described as a method where an investigator’s choice of a set of *n* objectives defines an *n*-dimensional volume in which all Pareto solutions to the problem are found. Next, *n* anchor points are identified. Anchor Points are feasible solutions, in the objective space, that correspond to the best possible values for respective individual objectives. The values of the anchor points are normalized to eliminate deficiencies associated with scales of individual objectives. The solution space volume is then reduced through the use of an *n*-dimensional “Utopia” hyperplane. The Utopia plane is defined such that it contains all *n* anchor points. Finally, a set of evenly distributed points on the Utopia hyperplane serve to constrain the all but one of the objectives under consideration. Solving for the optimal value of the lone objective at each one of these points will result in calculation of a Pareto solution. For a full mathematical description of NNC see the manuscripts by Dr. Achille Messac and coworkers[47, 113].

It is interesting to note that Shoval et al.[49] recently showed that best trade-off phenotypes for any organism are the weighted averages of archetypes. In the NNC method anchor points represent these archetypes. Results from Shoval et al. also indicate that experimentally observed phenotypes are contained within simple geometric shapes that are akin to the Utopia line, plane, or hyper-plane (depending on the dimension of MO analysis) in NNC – *i.e*., the geometric space defined by the anchor points.

### Analysis of growth-related pH changes in the medium

*Rhodopseduomonas palustris* CGA009 was grown in modified photosynthetic medium[114] with low phosphate under anaerobic conditions. To observe how pH of the growth media was affected by bacterial growth on various carbon sources, we lowered the phosphate concentration to 20% (5 mM) of the original concentration and adjusted the initial pH to 6.7 prior to cell inoculation. An organic source of acetate (10 mM), benzoate (3 mM), 4-hydroxylbenzoate (2.2 mM) or 4-coumarate (2 mM) was provided as the sole carbon source. Anaerobic cultures were placed 20 cm away from a 60 W incandescent light bulb under constant light, and optical density (OD at 660 nM) was monitored to calculate doubling time. The pH of the spent media was measured after cultures reached late exponential phase. Three biological replicates were included for each condition.

## List of abbreviations

4HBZ: 4-hydroxybenzoate
CBB: Calvin-Benson-Bassham
FBA: Flux balance analysis
GSM: Genome-scale model
LN: Light, anaerobic condition
MO: Multi-objective
MOFA: Multi-objective flux analysis
NNC: Normalized normal constraint
PHB: polyhydroxybutyrate
PNS: Purple non-sulfur
PO: Pareto-optimal
RP: *Rhodopseudomonas palustris*
rTCA: reductive citric acid cycle

## Declarations

### Ethics approval and consent to participate

Not applicable

### Consent for publication

Not applicable

### Availability of data

All data generated during this study are included in this published article (and its Supplementary Information files).

### Competing Interests

The authors declare that they have no competing interests.

### Funding

This research was supported by the LLNL Biofuels Scientific Focus Area, funded by the U.S. Department of Energy Office of Science, Office of Biological and Environmental Research Genomic Science program under FWP SCW1039 and Laboratory Research and Development program (14-ERD-091) at LLNL.

### Author’s contributions

AN developed the model, conducted all in silico analysis, and wrote the paper. YQ edited the paper and conducted all experimental analyses. SEW conduced the molecular dynamics simulations and edited the paper. JPR managed the project and edited the manuscript.

## Acknowledgments

The authors would like to thank Professor Jake McKinlay for sharing his time and expertise on *R. palustris* metabolism. We also would like to thank Dr. Peter Weber for his insightful comments and help editing this document. Work at LLNL was performed under the auspices of the U.S. Department of Energy by Lawrence Livermore National Laboratory under Contract DE-AC52–07NA27344. LLNL-JRNL-727615

